# Extensive recombination-driven coronavirus diversification expands the pool of potential pandemic pathogens

**DOI:** 10.1101/2021.02.03.429646

**Authors:** Stephen A. Goldstein, Joe Brown, Brent S. Pedersen, Aaron R. Quinlan, Nels C. Elde

## Abstract

The ongoing SARS-CoV-2 pandemic is the third zoonotic coronavirus identified in the last twenty years. Enzootic and epizootic coronaviruses of diverse lineages also pose a significant threat to livestock, as most recently observed for virulent strains of porcine epidemic diarrhea virus (PEDV) and swine acute diarrhea-associated coronavirus (SADS-CoV). Unique to RNA viruses, coronaviruses encode a proofreading exonuclease (ExoN) that lowers point mutation rates to increase the viability of large RNA virus genomes, which comes with the cost of limiting virus adaptation via point mutation. This limitation can be overcome by high rates of recombination that facilitate rapid increases in genetic diversification. To compare dynamics of recombination between related sequences, we developed an open-source computational workflow (IDPlot) to measure nucleotide identity, locate recombination breakpoints, and infer phylogenetic relationships. We analyzed recombination dynamics among three groups of coronaviruses with noteworthy impacts on human health and agriculture: *SARSr-CoV, Betacoronavirus-1*, and SADSr-CoV. We found that all three groups undergo recombination with highly diverged viruses from sparsely sampled or undescribed lineages, which can disrupt the inference of phylogenetic relationships. In most cases, no parental origin of recombinant regions could be found in genetic databases, suggesting that much coronavirus diversity remains unknown. These patterns of recombination expand the genetic pool that may contribute to future zoonotic events. Our results also illustrate the limitations of current sampling approaches for anticipating zoonotic threats to human and animal health.

## Introduction

In the 21^st^ century alone three zoonotic coronaviruses have caused widespread human infection: SARS-CoV in 2002 [1, 2], MERS-CoV in 2012 [2], and SARS-CoV-2 in 2019 [3]. Four other coronaviruses, OC43, 229E, NL63, and HKU1 are endemic in humans and cause mild-to-moderate respiratory disease with low fatality rates, though they may cause outbreaks of severe disease in vulnerable populations [4–7]. Like SARS-CoV-2, SARS-CoV, and MERS-CoV, these endemic viruses emerged from animal reservoirs. The origins of 229E and NL63 have been convincingly linked to bats, much like the 21^st^ century novel coronaviruses [8–10]. In a striking parallel, both MERS-CoV and 229E appear to have emerged from bats into camelids, established a new persistent reservoir, and then spilled over into humans [11–14]. In contrast, the viral lineages that include OC43 and HKU1 originated in rodents [15,16], though very limited rodent sampling leaves us with a poor understanding of the deep evolutionary history of these viruses. Given the short infectious period of human coronavirus infections, the establishment of endemicity was likely preceded by a period of intense and widespread transmission on regional or global scales. In other words, SARS-CoV-2 is likely the fifth widespread coronavirus epidemic or pandemic involving a still-circulating virus, though the severity of the previous four cannot be reliably ascertained.

Livestock are similarly impacted by spillover of coronaviruses from wildlife reservoirs. Three viruses closely related to OC43, bovine coronavirus (BCoV), equine coronavirus (ECoV) and porcine hemagglutinating encephalomyelitis virus (PHEV) are enzootic or epizootic in cows, horses, and pigs respectively [17]. Since 2017, newly emerged swine acute diarrhea syndrome-associated coronavirus (SADS-CoV) has caused significant mortality of piglets over the course of several outbreaks [18,19]. Sampling of bats proximal to impacted farms determined that SADS-CoV outbreaks are independent spillover events of SADSr(elated)-CoVs circulating in horseshoe bats [20]. Molecular studies of SADS-CoV have identified the potential for further cross-species transmission, including the ability to infect primary human airway and intestinal cells [21,22].

Emergence of novel viruses requires access to new hosts, often via ecological disruption, and the ability to efficiently infect these hosts, frequently driven by adaptive evolution. Uniquely among RNA viruses, coronavirus genomes encode a proofreading exonuclease that results in a significantly lower mutation rate for coronaviruses compared to other RNA viruses [23,24]. This mutational constraint is necessary for maintaining the stability of the large (27-32 kb) RNA genome but limits the evolution of coronaviruses via point mutation. The high recombination rate of coronaviruses compensates for the adaptive constraints imposed by high-fidelity genome replication [24,25]. The spike glycoprotein in particular has previously been identified as a recombination hotspot [26]. Acquisition of new spikes may broaden or alter receptor usage, enabling host-switches or expansion of host range. Additionally, it may result in evasion of population immunity within established host species, effectively expanding the pool of susceptible individuals. Recombination in other regions of the genome is less well-documented but may also influence host range, virulence, and tissue tropism, and likely contributed to the emergence of SARS-CoV [27,28].

To study the dynamics of recombination among clinically significant coronavirus lineages we developed a novel web-based software, IDPlot, that incorporates multiple analysis steps into a single user-friendly workflow. Analyses performed by IDPlot include multiple sequence alignment, nucleotide similarity analysis, and tree-based breakpoint prediction using the GARD algorithm from the HyPhy genetic analysis suite [29]. IDPlot also allows the direct export of sequence regions to NCBI Blast to ease identification of closest relatives to recombinant regions of interest.

Using IDPlot, we analyzed recombination events in three clinically significant lineages of coronaviruses with sufficient samplings to conduct robust analyses: SARS-CoV-2-like viruses, OC43-like viruses (*Betacoronavirus-1*) in the *Betacoronavirus* genus, and the SADSr-CoV group of alphacoronaviruses. In all three groups, we found clear evidence of recombination resulting in viruses with high overall nucleotide identity but exhibiting substantial genetic divergence in discrete genomic regions. Recombination was particularly enriched around and within the spike gene and 3’ accessory genes. Within all three groups, recombination has occurred with undescribed lineages, indicating that coronavirus diversity, even within these consequential sub-groups of viruses, remains considerably undersampled. The potential for viruses to rapidly acquire novel phenotypes through such recombination events underscores the importance of a more robust and coordinated ecological, public health, and research response to the ongoing pandemic threat of coronaviruses.

## Results

### Coronavirus phylogenetic relatedness is variable across genomes

Coronavirus genomes, at 27-32 kilobases (kb) in length, are among the largest known RNA genomes, surpassed only by invertebrate viruses in the same *Nidovirales* order [30,31]. The 5’ ~20 kb of the genome comprises open reading frames 1a and 1b, which are translated directly from the genome as polyproteins pp1a and pp1ab and proteolytically cleaved into constituent proteins (**Figure 1A**) [32]. Orf1ab is among the most conserved genes and encodes proteins essential for replication, including the RNA-dependent RNA-polymerase (RdRp), 3C-like protease (3ClPro), helicase, and methyltransferase. Given the high degree of conservation in this region, coronavirus species classification is typically determined by the relatedness of these key protein-coding regions [33]. The 3’ ~10 kb of the genome contains structural genes including those encoding the spike and the nucleocapsid proteins, as well as numbered accessory genes that are unique to coronavirus genera and subgenera [34]. In contrast to the relative stability of the replicase region of the genome, the structural and accessory region, and in particular the spike glycoprotein, have been identified as recombination hotspots [26].

**Figure 1.**
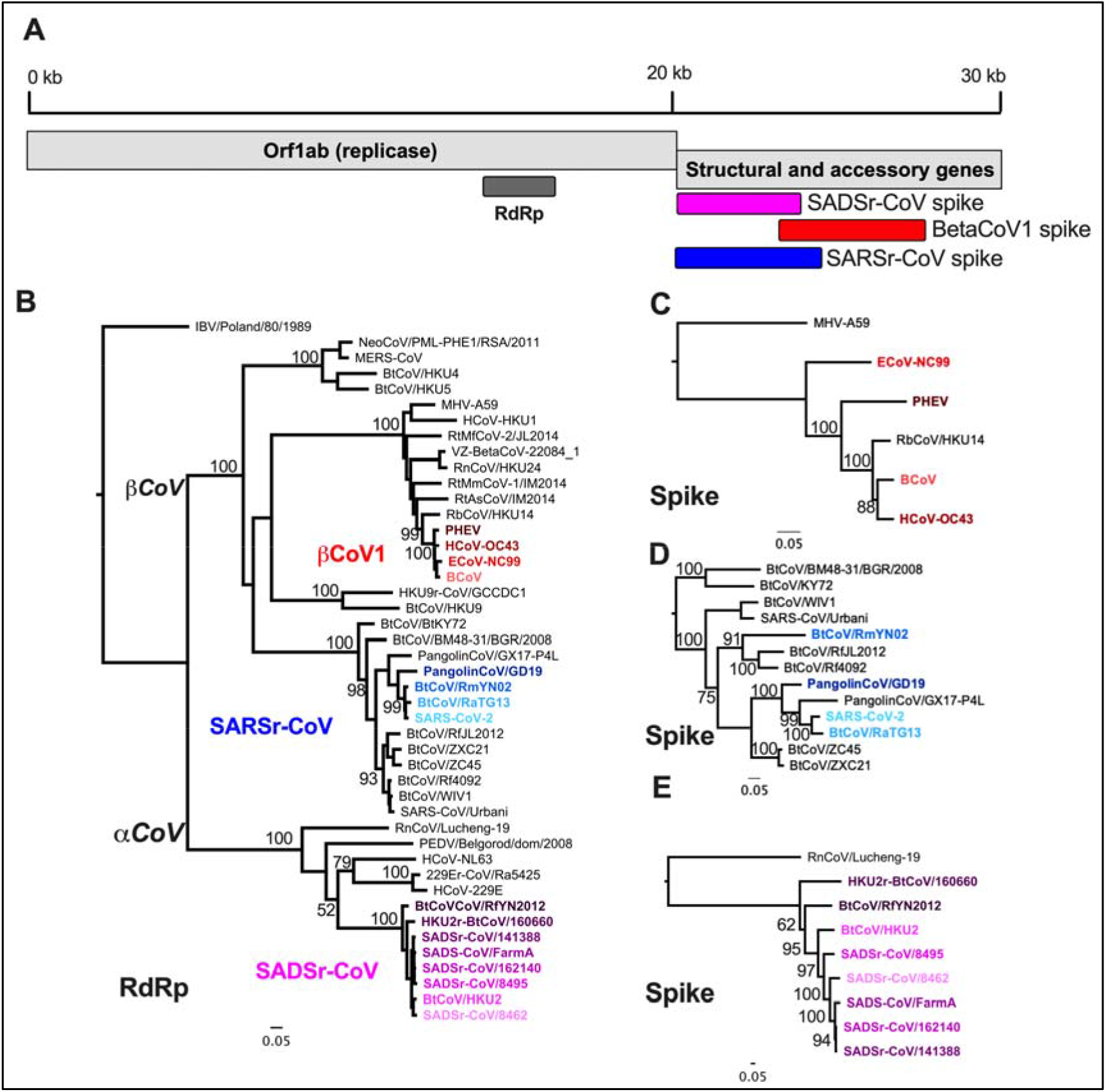
AlphaCoV and BetaCoV phylogenetic relationships are genome region-dependent. A) Basic coronavirus genome organization with the 5’ ~20 kb comprising the replicase gene that is proteolytically processed into up to 16 individual proteins. The 3’ 10 kb comprises structural and genus-specific accessory genes. B) Maximum-likelihood (ML) phylogenetic tree of alpha and betaCoVs fulllength RNA-dependent RNA-polymerase encoding region of Orf1ab. C) ML phylogenetic tree of fulllength spike genes from SADS-related CoVs (magenta), rooted with the distantly related alphacoronavirus HCoV-229E D) ML phylogenetic tree of spike genes from viruses in the species *Betacoronavirus 1* (red) rooted with the distantly related betacoronavirus mouse hepatitis virus. E) ML phylogenetic tree of spike genes of SARSr-CoVs, with SARS-CoV-2-like viruses further analyzed in the paper highlighted in blue.

We set out to characterize the role of recombination in generating diversity across the coronavirus phylogeny. A classic signature of recombination is differing topology and/or branch lengths of phylogenetic trees depending on what genomic regions are analyzed. To identify lineages of interest for recombination analysis, we built a maximum-likelihood phylogenetic tree of full-length RdRp-encoding regions of representative alpha and betacoronaviruses, which contain all human and most mammalian coronaviruses (**Figure 1B**). To further test whether comparisons of RdRp sequence reflected ancestral relatedness, we conducted the same analysis for the 3ClPro and Helicase-encoding regions of Orf1ab (**Figure S2**). Phylogenetic relationships were generally maintained in these trees and genetic relatedness remains very high (90-99% within groups), which leads to some reshuffling with low bootstrap support. From these trees we chose to further investigate the evolutionary dynamics of three clinically significant groups of coronaviruses: SARS-CoV-2 like viruses (blue) from within *SARSr(elated)-CoV*, among which recombination has been reported though not characterized in detail, endemic and enzootic OC43-like viruses of *Betacoronavirus-1* (*BetaCoV1*) (red), and SADSr-CoVs (magenta). Although other coronavirus lineages are of public health interest, such as those including the human coronaviruses HKU1, NL63, and 229E there is a relative paucity of closely related sequences to these viruses, limiting our current ability to analyze these genomes.

Within each group there is modest diversity revealed by comparing RdRp sequence: 94-99% nt identity among the SADSr-CoVs, >97% nt identity within *Betacoronavirus-1*, and 91-99% among the SARS-CoV-2-like viruses (**Figure S3**). Similar results were observed for 3ClPro and Helicase-encoding regions (**Figure S4**). In contrast, spike gene phylogenetic trees of each group show greater diversity as illustrated by extended branch lengths and/or changes in tree topology, suggesting either rapid evolution and/or recombination diversifies this region (**Figure 1B-D**). To analyze these possibilities, we developed a new pipeline to better study these evolutionary patterns.

### IDPlot Facilitates Nucleotide Identity and Recombination Analysis

To investigate possible recombination-driven diversity among these viruses we developed IDPlot, which incorporates several distinct analysis steps into a single Nextflow workflow [35] and generates a comprehensive HTML report to facilitate interpretation and downstream analysis. IDPlot combines existing algorithms into a single pipeline and provides a statistically supported means to adjust recombination prediction and phylogenetic analysis in a quickly interpretable visual display.

First, IDPlot generates a multiple sequence alignment using MAFFT (**Figure 2A**) [36] with user-assigned reference and query sequences. In its default configuration, IDPlot then generates a sliding window average nucleotide identity (ANI) plot, also displaying the multiple sequence alignment with differences to the reference sequence (colored vertical lines) and gaps (gray boxes) clearly highlighted. The plot is zoomable, and selected sequence regions can be exported directly to NCBI BLAST. Users can also choose to run GARD, the recombination detection program from the HyPhy suite of genomic analysis tools [29]. If GARD is implemented (**Figure 2B**), distinct regions of the multiple sequence alignment are depicted between the alignment and the ANI plot, and phylogenetic trees for each region are generated using FastTree2 (**Figure 2C**) [37] and displayed (**Figure 2E**).

**Figure 2.**
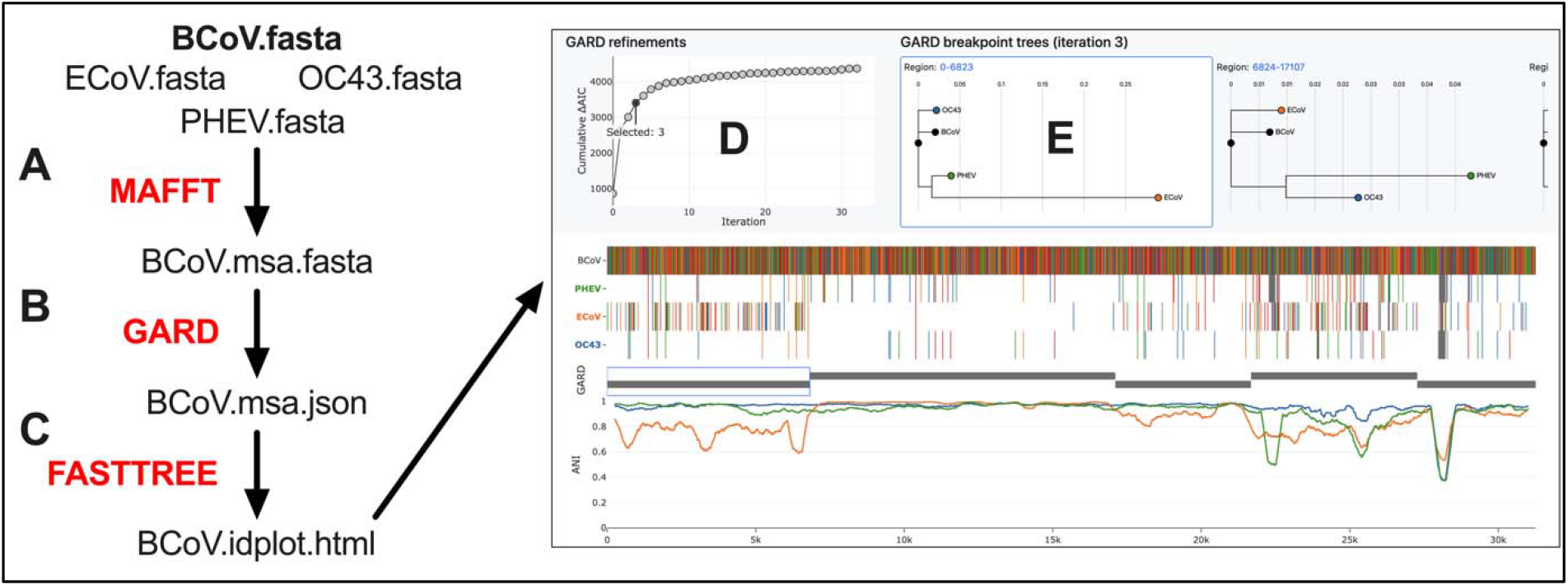
IDPlot workflow. IDPlot workflow. A) Reference and query sequences are aligned using MAFFT. B) Breakpoint detection is performed using GARD, capturing breakpoints across iterative refinements. C) Phylogenetic trees based on breakpoints from each iteration and are created using FastTree 2. D) Improvement in ΔAIC-c is plotted against the iteration E) Phylogenetic trees associated with the selected GARD iteration are displayed

A significant barrier to effective use of GARD is that because it ultimately presents multiple (sometimes dozens of) iterations, model choice and therefore the selection of breakpoints for further analysis can be challenging. To alleviate this issue the IDPlot output includes a graph showing a cumulative count of GARD’s statistical iterations (Akaike information criterion (AIC-c) on the y-axis) and the GARD model number on the x-axis (**Figure 2D**). GARD uses ΔAIC-c for each proposed model to indicate the degree of fit improvement over the preceding model, and this graph allows the user to easily determine when improvements become increasingly marginal, which is often accompanied by prediction of spurious breakpoints. Upon selection of a GARD iteration, the display switches to show the associated phylogenetic trees (**Figure 2E**). Genomic regions are clickable, immediately bringing the appropriate phylogenetic tree to the center of the display. Finally, the ability to export sequences directly to BLAST enables the user to search for related sequences in GenBank, useful when defined regions are highly divergent from the reference sequence or others included in the data set under analysis.

#### SARS-CoV-2-like virus recombination with distant SARSr-CoVs

To test and validate IDPlot as a tool for examining the recombination dynamics of coronaviruses, we initially conducted an analysis of SARS-CoV-2-like viruses within *SARSr-CoV*. We chose these viruses as our initial IDPlot case study because recombination has been previously described [38,39], though not characterized in great phylogenetic detail. This provided the opportunity to evaluate IDPlot against a known framework but also advance our understanding of the role recombination has played in the evolution of these clinically significant viruses.

Prior to 2019 the SARS-CoV-2 branch within *SARSr-CoV* was known only from a single, partial RdRp sequence published in 2016 [40]. Upon the discovery of SARS-CoV-2 this RdRp sequence was extended to full genome-scale [3] and additional representatives from bats and pangolins have since been identified [39] [41,42]. However, this singularly consequential lineage remains only sparsely sampled and its evolutionary history largely obscured. Most attention on these viruses to date has focused on the recent evolutionary history of SARS-CoV-2 with respect to possible animal reservoirs and recombinant origins. Much less attention has been paid to analyzing the evolution of known close relatives, the bat viruses RaTG13 and RmYN02, and PangolinCoV/GD19.

Our IDPlot analysis does not support an emergence of SARS-CoV-2 via recent recombination, consistent with previously published work [3] [38]. RaTG13 shows consistently high identity across the genome with the only notable dip comprising the receptor-binding domain in the C-terminal region of spike S1 (**Figure 3A**), which is proposed to originate via either recombination or diversifying selection [38]. However, the still limited sampling in the SARS-CoV-2-like lineage results in weak phylogenetic signals unable to distinguish between rapid mutational divergence and recombination producing the low ANI in the RaTG13 receptor binding domain.

**Figure 3.**
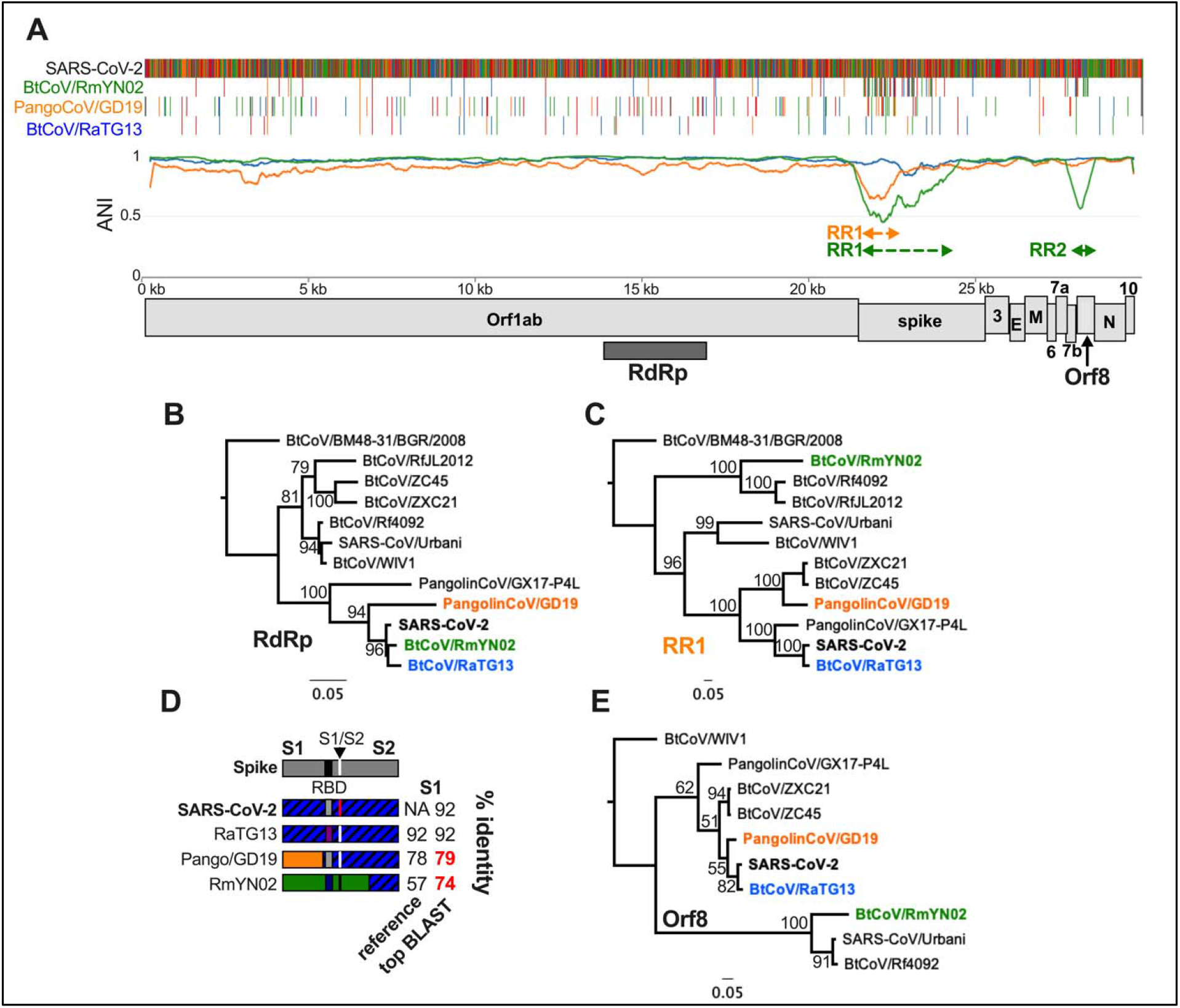
SARSr-CoV IDPlot Analysis. A) IDPlot analysis of SARS-CoV-2-like SARSr-CoVs with color-coded dashed lines defining divergent regions arising from recombination events with ancestral viruses. B) ML tree of the RdRp-encoding region of SARS-2-like and other SARSr-CoVs showing close relationship between the SARS-CoV-2-like viruses. C) ML tree of PangolinCoV/GD19 RR1 (which overlaps with BtCoV/RmYN02 RR1) showing different topology than the RdRp tree. D) Schematic of spike proteins indicating divergent regions and nucleotide identity to the reference sequence and closest related sequence in GenBank. E) ML tree of ORf8 showing that RmYN02 Orf8 is a divergent member of the SARS-CoV-like Orf8 branch.

In contrast, PangolinCoV/GD19 and RmYN02 show one and two significant drops in ANI, respectively. Phylogenetic analysis of the PangolinCoV/GD19 recombinant region captures the signal for both that virus (**Figure 3A, 3C, S5C**) and RmYN02 RR1, showing that both viruses fall onto separate branches highly divergent from SARS-CoV-2 and RaTG13 (**Figures 3C**) with only 81% and 74% nucleotide identity to the closest sequences in GenBank, respectively (**Figure 3D, S5A**). These findings identify three unique spike genes among SARS-CoV-2 and its three closest known relatives (**Figure 3D**), indicative of recombination with *SARSr-CoV* lineages that remain to be discovered despite being the focus of intense virus sampling efforts over the last eighteen years, since the emergence of SARS-CoV.

In addition to spike, RmYN02 contains a second recombinant region that encompasses the 3’ end of Orf7b and the large majority of Orf8 (**Figure 3A, S5A**). Orf8 is known to be highly dynamic in SARSr-CoVs. SARS-CoV underwent an attenuating 29 nt deletion in Orf8 in 2002-2003 [43] and Orf8 deletions have been identified in numerous SARS-CoV-2 isolates as well [44–46]. In bat SARSr-CoVs intact Orf8 is typically though not always present but exhibits a high degree of phylogenetic incongruence. Additionally, the progenitor of SARS-CoV encoded an Orf8 gene gained by recombination [28,47]. The BtCoV/RmYN02 Orf8 has only 50% nt identity to SARS-CoV-2 Orf8 and groups as a distantly related member of the branch containing SARS-CoV (**Figure 3E**), exhibiting just 80% nucleotide identity to the closest known sequence. Although the precise function of Orf8 is unknown, there is some evidence that like other accessory proteins it mediates immune evasion [43]. Therefore, recombination in Orf8 has the potential to alter virus-host interactions and may, like spike recombination, impact host range and virulence.

This analysis confirmed that IDPlot allows us to characterize recombination events in detail with a single workflow. We demonstrate that multiple SARS-CoV-2-like viruses have recombined with unsampled SARSr-CoV lineages, limiting our ability to assess sources of genetic diversification for these viruses. Under-sampling has implications limiting the incisiveness of both laboratory and field investigations of these viruses.

### OC43-like viruses encode divergent spikes acquired from unsampled betacoronaviruses

After validating IDPlot for recombination analysis of coronaviruses, we used it to characterize recombination among the viruses in the *Betacoronavirus-1* (*BetaCov1*) group, which includes the human endemic coronavirus OC43 and closely related livestock pathogens bovine coronavirus (BCoV), equine coronavirus (ECoV), porcine hemagglutinating encephalomyelitis virus (PHEV), and Dromedary camel coronavirus HKU23 (HKU23). Due to the apparent low virulence of OC43 and limited sampling of the lineage, these viruses receive relatively little attention outside agricultural research. However, this lineage has produced a highly transmissible human virus, can cause severe disease in vulnerable adults, and is poorly sampled [4]. An ancestral BCoV is believed to be the progenitor of the other currently recognized *BetaCoV1* viruses with divergence dates estimated at 100-150 years ago for OC43/PHEV [48] and 50 years ago for HKU23 [49]. Recombination with other betacoronaviruses has been previously described for HKU23, so we excluded it from our analysis [50]. The most closely related known virus to *BetaCoV1*, rabbit coronavirus HKU14 (RbCoV/HKU14) was reported to associate with ECoV in some regions [51], but no detailed recombination analysis of the relationship between these viruses has been previously described.

We conducted IDPlot analysis of OC43 and these related enzootic viruses of livestock (**Figure 4A**) and identified at least six major recombination breakpoints in the ECoV genome. The largest divergent region (Region 2) is >6 kilobases (**Figure 4A**). This region encompassing ~20% of the genome exhibits only ~75% nt identity to the reference sequence, just ~81% identity to any known sequence, and occupies a distant phylogenetic position relative to RdRp (**Figure 4B-C, S6A, S6C-D**). In contrast to previous reports that ECoV clusters closely with RbCoV/HKU14 in this region [51], our analysis reveals that this region of ECoV was acquired via recombination from a viral lineage not documented in GenBank.

**Figure 4.**
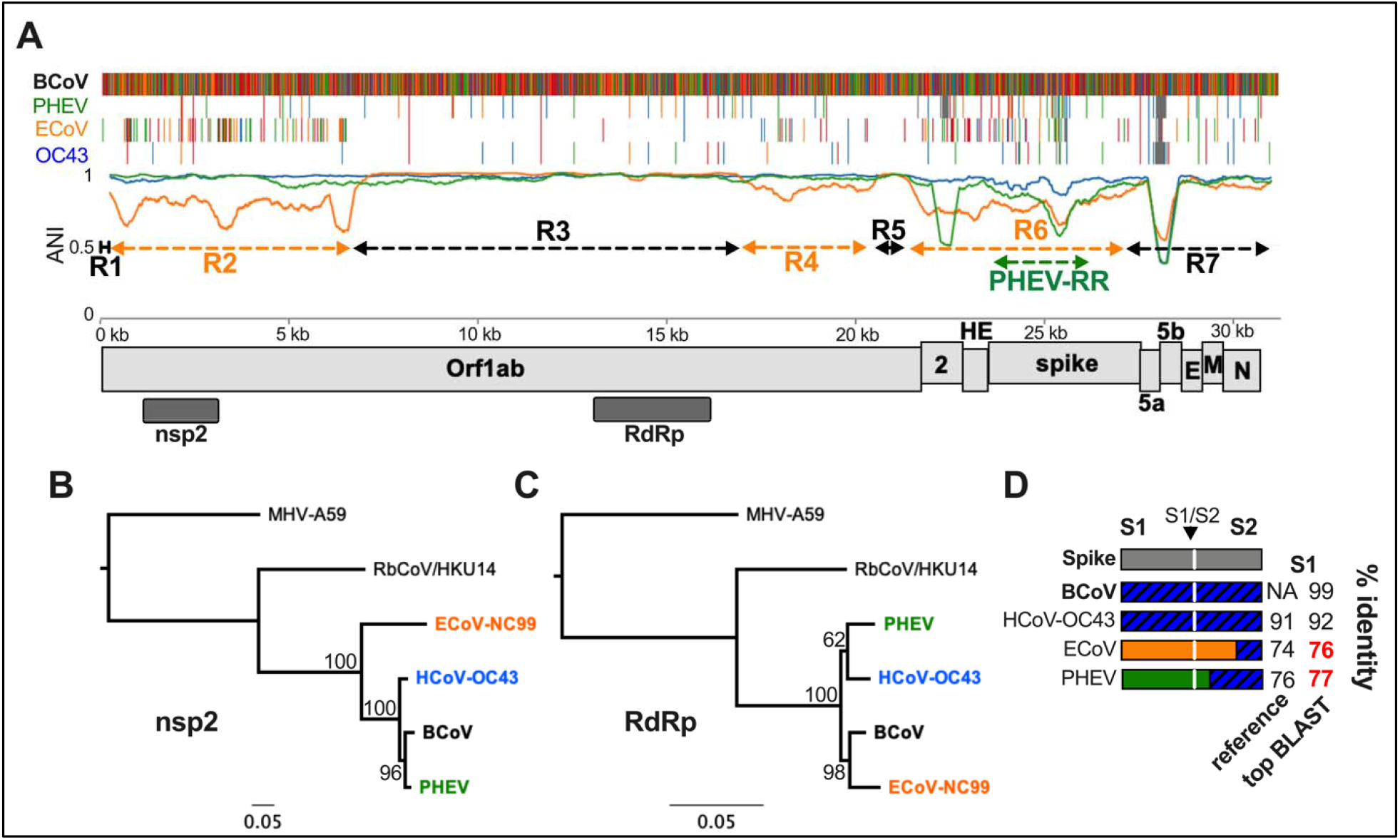
Recombination analysis of *Betacoronavirus-1*. A) Nucleotide identity plot and multiple sequence alignment of *BetaCoV-1* viruses. Orange dashed lines indicate divergent regions of the ECoV-NC99 genome while black dashed lines are regions with high identity to the reference sequence bovine coronavirus (BCoV). B) ML tree of nsp2-encoding region of Orf1ab, which falls within the divergent ECoV-NC99 Region 2. C) ML tree of the RdRp-encoding region of Orf1ab. D) Schematic depicting the spike gene diversity of *BetaCoV1* demonstrating the divergence of ECoV-NC99 and PHEV. Top BLAST hits in bolded red indicate no GenBank entries with >80% nucleotide identity.

Striking variability in ANI within Region 2 led us to conduct a more detailed analysis. IDPlot did not predict internal Region 2 breakpoints, so we conducted a manual analysis guided by the IDPlot multiple sequence alignment, phylogenetic trees for each proposed sub-region, and BLAST analysis to further dissect differing evolutionary relationships for sub-regions. We found at least six and possibly seven distinct sub-regions (**Figure S7**). Nucleotide identity to top BLAST hits of these sub-regions is highly variable (<70% to >90%), as is identity of the hits themselves, with genetic contribution from RbCoV/HKU14-like viruses, BCoV-like viruses, and more distant, uncharacterized lineages within the *Embecovirus* genus (**Figure S5**). Together, this demonstrates that Region 2 was not acquired via a single recombination event but rather represents a mosaic of known and unknown viral lineages that share an overlapping ecological niche with ancestral ECoV.

Another major recombinant ECoV region, Region 6, includes the entire NS2 and HE genes as well as the majority of the spike gene (**Figure 4A, S6A**). Within this region on the multiple sequence alignment, we also identified a recombination event encompassing the majority of the PHEV spike gene, though this required downsampling (removing ECoV) to simplify the analysis (**Figure 4A, S6A)**. Both ECoV Region 6 and the PHEV recombinant region occupy relatively distant nodes on a phylogenetic tree (**Figure S6G, J**) and exhibit <80% sequence identity to the reference sequence or any sequence in GenBank (**Figure S6A**), indicating they are derived from independent recombination events or diverged via repeated mutations. Recombination appears most likely given the even dispersion of low identity throughout the region, including in portions of the spike S2 domain which is otherwise highly conserved. Additional sampling might more definitively resolve these possibilities. Finally, we identified a third recombinant region, Region 4, in which ECoV exhibited high nucleotide identity with RbCoV/HKU14 (**Figure S4A, E**), further demonstrating the highly mosaic nature of the ECoV genome.

Our analysis of equine coronavirus offers a remarkable example of the degree and speed of divergence facilitated by the high recombination rates among coronaviruses. Previous genomic characterization of ECoV suggested that it is the most divergent member of *BetaCoV1* based on nucleotide identity and phylogenetic positioning of full-length Orf1ab. However, in the >10 kilobase Region 3 that accounts for ~1/3 of the entire genome (**Figure 4A**) ECoV exhibits the highest nucleotide identity to BCoV in our dataset (98.5%) (**Figure 4A, S6D**), which is inconsistent with it having diverged earlier than OC43 and PHEV. The latter viruses are estimated to have shared a common ancestor with BCoV 100-150 years ago [48], suggesting that all of the observed ECoV recombination has occurred more recently. Our discovery of recombinant regions of unknown betacoronavirus origin suggest that unsampled, distantly related lineages occupy overlapping ecological niches with ECoV and may continue to circulate and participate in recombination events. Basal members of the subgenus that includes *BetaCoV1* have been identified exclusively in rodents (**Figure 1B**), suggesting they are a natural reservoir for these viruses. Although relatively little attention has been directed to these viruses, studies of BCoV and ECoV cross-neutralization suggest population immunity to OC43 may provide only limited protection against infection mediated by these novel spikes [52]. No recent zoonotic infections from this lineage have been documented, but the genomic collision of these viruses with yet-undiscovered, presumably <u>rodent</u> viruses warrants a reassessment of their potential threat to human health.

#### SADSr-CoVs encode highly diverse spike and accessory genes

In 2017 a series of highly lethal diarrheal disease outbreaks on Chinese pig farms were linked to a novel alphacoronavirus, swine acute diarrhea syndrome-associated coronavirus (SADS-CoV) [20,53], which is closely related to the previously described BtCoV/HKU2 [54]. Sampling of horseshoe bats nearby affected farms revealed numerous SADSr-CoVs with >95% genome-wide nucleotide identity, suggesting porcine outbreaks were due to spillover from local bat populations. To gain a better view of the genetic diversity among these viruses, we conducted IDPlot analysis of a prototypical SADS-CoV isolate (FarmA) and seven bat SADSr-CoVs sampled at different times before and after the first outbreaks in livestock (**Figure 5A**) using bat SADSr-CoV/162140 as a reference sequence. Three notable observations emerged from the identity plot: 1. Like ECoV, BtCoV/RfYN2012 exhibits evidence of recombination in the 5’ end of Orf1ab 2. the spike region of the genome is highly variable as previously reported [20], and 3. The 3’ end of the genome also exhibits considerable diversity (**Figure 5A**).

**Figure 5.**
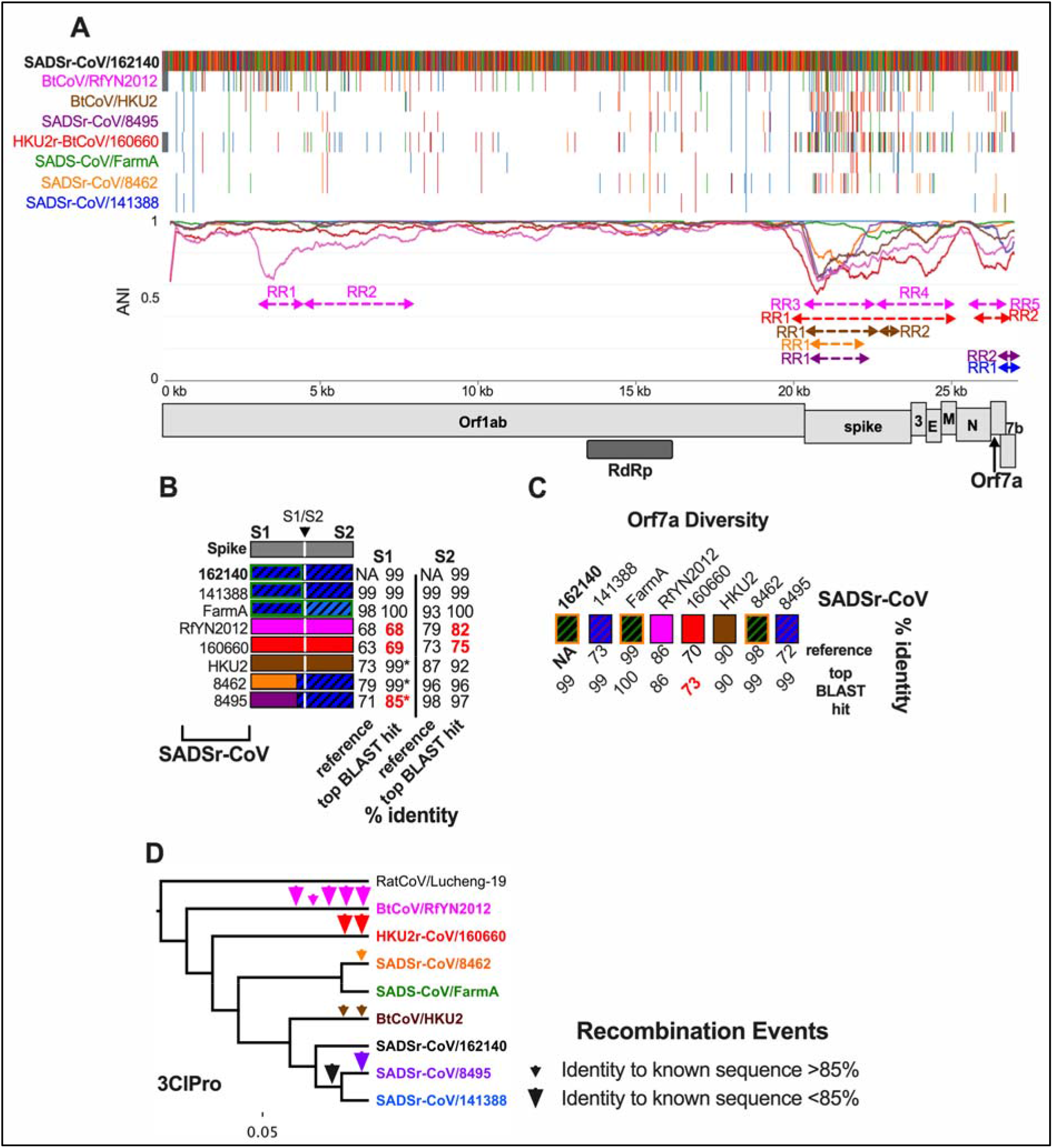
SADSr-CoV IDPlot Analysis. A) IDPlot nucleotide identity and multiple sequence alignment of eight SADSr-CoVs. Color-coded dashed lines indicate divergent regions in corresponding viruses owing to recombination events. B) Schematic of spike genes of SADSr-CoVs along with nucleotide identity to the reference sequence and closest related sequences in GenBank for S1 and S2 domains. C) Schematic of Orf7a diversity with nucleotide identity to the reference sequence and closest related sequences in GenBank D) Phylogenetic tree of SADSr-CoVs based on 3ClPro sequence illustrating the history of inferred recombination events indicated by arrowheads.

To confirm the recent common ancestry of SADSr-CoVs in our data set we conducted nucleotide identity and phylogenetic analyses of the RdRp, 3ClPro, helicase, and methyltransferase NTD-encoding regions of Orf1ab. All viruses exhibit exhibit 94-100% nucleotide identity to the reference SADSr-CoV/162140 in these regions of the genome (**Figure S4E-F, S8B-E, S9B-C**). In contrast, BtCoV/RfYN2012 recombinant region 1 (RR1) has <70% identity to the reference or any known sequence (**Figure S8F, S9A**), providing evidence that an uncharacterized alphacoronavirus lineage circulates in horseshoe bats, which frequently recombines with SADSr-CoVs.

The spike gene is a striking recombination hotspot among SADSr-CoVs. Due to the clustering of putative breakpoints surrounding the 5’ end, 3’ end, and middle of spike, we ran IDPlot on subsets of three viruses – SADSr-CoV/162140 (reference), SADSr-CoV/141388 or SADS-CoV/FarmA, and a virus of interest from the larger dataset. We found breakpoints delineating six distinct and highly divergent spike genes among the eight analyzed viruses (**Figure 5B**), which reflects recombination events encompassing either the entire spike or the S1 subunit that mediates receptor binding. There are 3 unique full-length spikes (BtCoV/RfY2012, HKU2r-BtCoV/160660, BtCoV/HKU2) with 63-73% nucleotide identity to the reference sequence and two unique S1 domains (SADSr-CoVs/8462 and 8495) with <80% identity to the reference (**Figure 5B, S9A**). Some of these regions match with high identity to partial sequences in GenBank (indicated by an asterisk in Figure 5B) which may be either the parent virus of the recombinant spike or different isolates of the same virus for which a full-length genome is available. Other spikes in this dataset are clearly divergent from any other known sequence.

In addition to spike, accessory proteins that target innate immunity can play important roles in host range and pathogenesis [34]. We found a second recombination hotspot surrounding the accessory gene Orf7a, which rivals spike gene diversification. Specifically, our dataset contained five distinct Orf7a genes, some of which lack any closely related sequences in GenBank (**Figure 5C, S9A**).

Finally, we mapped each inferred occurrence of a recombination event onto a SADSr-CoV phylogenetic tree. SADSr-CoVs 141388 and 8495 share an Orf7a recombination event, suggesting a recent common ancestor for these two viruses. The tree based on 3ClPro was most consistent with this evolutionary scenario (**Figure 5D**), while the other trees exhibit slightly different topology with minimal diversity, likely due to cryptic recombination events among very closely related viruses. Considering the 3ClPro tree, it is evident that many independent recombination events occurred in the very recent past given that few of the events are shared among the viruses in our dataset (**Figure 5D**).

The SADSr-CoV lineage is rapidly diversifying via recombination, particularly in the spike and ORF7a accessory genes. We observed that numerous viruses with >95-99% identity in conserved Orf1ab regions contain highly divergent spike and accessory genes which may shift host range and virulence in otherwise nearly isogenic viruses. These findings highlight how viruses sampled to date represent only a sliver of circulating SADSr-CoV coronavirus diversity and that coronaviruses can change rapidly, drastically, and unpredictably via recombination with both known and unknown lineages. The SADSr-CoVs exemplify the potential of coronaviruses to rapidly evolve through promiscuous recombination.

## Discussion

We developed IDPlot to explore the role of recombination in the diversification of coronaviruses. Coronaviruses are ubiquitous human pathogens with vast and underexplored genetic diversity. SARS-CoV-2 is the second SARSr-CoV known to infect humans and the fifth zoonotic coronavirus known to sweep through the human population following HCoVs 229E, NL63, HKU1, and OC43 [9,10,15,48,55,56]. Most effort in evaluating the threat to human health posed by coronaviruses has been dedicated to discovery of novel SARSr-CoVs in wildlife, yet prior to the SARS-CoV-2 pandemic this group of viruses went largely undetected. Much less attention has been paid to other groups that have produced human coronaviruses such as the sparsely sampled *Betacoronavirus-1* and emerging livestock viruses such as the SADSr-CoVs, which exhibit potential to infect humans and already have significant economic impacts. Recombination detection can be difficult when parental viruses are unknown, as was revealed with our analysis, due to difficulty in distinguishing between true recombination events versus repeated mutations under strong selective pressure. Rapid evolution is most evident for spike receptor binding domains, leading to polymorphism at critical residues [57,58]. Multiple sequence alignments generated by IDPlot demonstrate that even in divergent spikes, the low nucleotide identity is evenly distributed throughout putative recombinant regions. Additionally, we see high divergence even in conserved regions such as Orf1ab and the spike S2 domain. In all of these regions, including accessory genes, reshuffling of phylogenetic trees described in our analysis provides strong evidence that recombination, not repeated individual mutations of critical amino acid residues, accounts for the observed diversity.

We initially used the SARS-CoV-2-like viruses to test and validate IDPlot and in the process characterized recombination among these viruses in greater detail than previously reported. The observed variability in arrangements of PangolinCoV/GD19 and RmYN02 on a *SARSr-CoV* phylogenetic tree (**Figure 3B-C, 3E, S5**) depending on the region being sampled is a classic recombination signal easily observed in the IDPlot output. We also analyzed recombination dynamics for viruses in *BetaCoV1* and among SADSr-CoVs. Broad similarities emerge from these studies. Most recombination appears to involve the spike gene and/or various accessory genes. However, in both *BetaCoV1* and among SADSr-CoVs we detected recombination events in Orf1ab as well. Spike and accessory gene recombination events are particularly notable given the potential to influence host range and pathogenesis.

This preliminary analysis showed that IDPlot is a powerful new pipeline for sequence identity analysis, breakpoint prediction, and phylogenetic analysis. Existing workflows for nucleotide similarity analysis are proprietary, lack the ability to identify phylogenetic incongruence that is a signature of recombination and do not support direct export of genomic regions for BLAST analysis. This automates and streamlines multi-step analysis with few barriers to use. Nevertheless, there are opportunities for further improvement. Analysis of recombination breakpoints implemented in GARD are of limited value for resolving unique breakpoints in close proximity, as observed surrounding and within SADSr-CoV and other spike genes, necessitating the use of small sets of sequences. Second, GARD is computationally intensive and best suited to small data sets. It is configured as an optional step in IDPlot, so multiple sequence alignments and nucleotide identity plots can be rapidly generated in a local environment. However, for GARD analysis we relied on a high-performance computing cluster to expedite the process. In the future, we anticipate adding other, less intensive breakpoint prediction algorithms to the IDPlot options menu. Future advances in computational methods may also improve the ability to resolve unique breakpoints clustered in genomic regions that are recombination hotspots, most notably the spike gene.

Our IDPlot analyses revealed new evidence of extensive recombination-driven evolution in other coronavirus groups. Wildlife sampling indicates that SADSr-Covs are a large pool of closely related viruses circulating in horseshoe bat populations at high frequency. This is the same genus of bats that include SARSr-CoVs suggesting that the ecological conditions for SADSr-CoV spillover into humans may be in place. The relatedness of these viruses means they have had little time to diverge via mutation, but we find they are rapidly diversifying due to recombination, acquiring spike and accessory genes from unsampled viral lineages. These findings demonstrate that rather than a single threat to human health posed by SADS-CoV, there is a highly diverse reservoir of such viruses in an ecological position and with diversity reminiscent of SARSr-CoVs. We found a similar dynamic at play among *BetaCoV1* which are under-sampled to an even greater degree and receive far less attention. Nevertheless, these viruses are involved in genetic exchange with unsampled lineages, with unpredictable consequences.

Our findings bear on strategies for anticipating and countering future zoonotic events. SARSr-CoVs garner considerable attention, with an intense focus on viruses able to infect human cells using ACE-2 as an entry receptor. However, RmYN02 demonstrates that viruses can toggle between spikes that recognize ACE-2 or different entry receptors but still infect the same hosts and continue to undergo recombination. Work to prepare for future zoonotic SARSr-CoVs must account for the possibility that the threat will come from coronaviruses only distantly related to SARSr-CoVs undergoing frequent recombination and distributing genetic diversity across the phylogenetic tree of coronaviruses.

More attention to the evolutionary dynamics of *BetaCoV1* and SADSr-CoVs is also warranted. Both groups originate in wildlife: rodents and horseshoe bats respectively, and are enzootic or epizootic in livestock. *BetaCoV1* includes a pandemic virus that swept the human population, OC43, while SADS-CoV efficiently infects primary human respiratory and intestinal epithelial cells [22]. Our ability to anticipate threats from both groups would benefit from additional sampling, with *BetaCoV1* being particularly undersampled. Increased surveillance at wildlife-livestock interfaces, including agricultural workers is needed for early detection of novel viruses coming into contact with humans. Due to recombination, prior infection with a virus such as OC43 cannot be presumed to be protective against even closely related viruses that can encode highly divergent spikes, as demonstrated in our analysis. Similarly, efforts to develop medical countermeasures against SADS-CoV should consider the full breadth of diversity among related viruses, while aiming for broadly effective vaccines and therapeutics.

Using IDPlot, we identified extensive diversity among coronavirus spike and accessory genes with potential implications for future pandemics. From the standpoint of understanding coronavirus evolution, frequent recombination events often reshuffle phylogenetic trees and can obscure evolutionary relationships. The extent to which viruses in current databases contain genomic regions with no known close relatives makes clear that coronavirus diversity is vast and poorly sampled, even for viruses circulating in well-studied locations. This proximity raises the possibility of recurrent zoonoses of coronaviruses encoding divergent spike and accessory genes. Therefore, preparedness efforts should consider a broad range of virus diversity rather than risk a more narrow focus on close relatives of coronaviruses that most recently impacted human health.

## Methods

### Virus Sequences

All sequences were downloaded from GenBank with the exception of PangolinCoV/GD19 and BtCoV/RmYN02, which were acquired from the Global Initiative on Sharing All Influenza Data (GISAID) database (https://www.gisaid.org).

### IDPlot

IDPlot is initiated by the user designating reference and query sequences. A .gff3 annotation file can also be included in the input. The first step of IDPlot is multiple sequence alignment using MAFFT [36] with default parameters. Size of the sliding window is customizable and set to 500 for all of our analyses. For recombination analysis we ran GARD [29] as an optional step, utilizing the multiple sequence alignment generated by MAFFT. Trees for each GARD iteration are generated and displayed using Fast Tree 2 [37]. The entire output is then exported into a chosen directory as idplot.html as well .json files containing raw GARD data. More detailed information on IDPlot is available in the GitHub repository at https://github.com/brwnj/idplot.

### Phylogenetic validation of breakpoints

Putative breakpoints were further tested by maximum-likelihood phylogenetic analysis using PhyML [59]. For *Betacoronavirus-1*, RbCoV/HKU14 and MHV (as a root) were aligned with the four viruses in the IDPlot dataset. For SADSr-CoVs we chose HCoV-229E as the root, with the exception of the spike gene, and aligned it with the eight viruses in our dataset. We rooted the SARSr-CoVs with BtCoV/BM48-31/BGR/2008. Given the better sampling of SARSr-CoV, we included more diversity in that alignment to enhance phylogenetic signal. The signal for *BetaCoV1* and SADSr-CoV is constrained by sampling limitations. We extracted breakpoint-defined regions from the alignment and generated ML-phylogenetic trees using a GTR substitution model and 100 bootstraps. “Up” and “Dn” regions are the 500 nucleotides upstream or downstream of a proposed 5’ or 3’ breakpoint, respectively. In the case of SADSr-CoV the clustering of breakpoints around the 5’ and 3’ ends of spike precluded using unique Up and Dn regions for each recombination event. Instead, we used the N-terminal section of nsp16 (MTase) and the M gene, respectively. For BtCoV/RmYN02 RR2 and ORf8 phylogenetic testing we excluded SARSr-CoVs that have a deletion in Orf8. RmYN02 UpRR2 also does not include BtCoV/WIV1 because it has a unique open reading frameed insert in this region and so does not align with SARSr-CoVs lacking this Orfx.

### BLAST analysis

To identify the source of recombinant regions we used NCBI Blastn with default parameters, excluding the query sequence from the search. For SADSr-CoVs partial spike sequences frequently appear as top hits. We included these, denoted by an asterisk in reporting the results.

## Acknowledgements

We thank E.C. Holmes and co-authors for the use of BtCov/RmYN02 and PangolinCoV/GD19 genome sequences in our analysis. We thank Z.A. Hilbert for manuscript assistance.

## References

1. Drosten C, Günther S, Preiser W, van der Werf S, Brodt H-R, Becker S, et al. Identification of a Novel Coronavirus in Patients with Severe Acute Respiratory Syndrome. The New England Journal of Medicine. 2003;348: 1967–1976. doi:10.1056/NEJMoa030747

2. Zaki AM, van Boheemen S, Bestebroer TM, Osterhaus ADME, Fouchier RAM. Isolation of a Novel Coronavirus from a Man with Pneumonia in Saudi Arabia. The New England Journal of Medicine. 2012;367: 1814–1820. doi:10.1056/NEJMoa1211721

3. Zhou P, Yang X-L, Wang X-G, Ben Hu, Zhang L, Zhang W, et al. A pneumonia outbreak associated with a new coronavirus of probable bat origin. Nature. Nature Publishing Group; 2020;579: 270–273. doi:10.1038/s41586-020-2012-7

4. Patrick DM, Petric M, Skowronski DM, Guasparini R, Booth TF, Krajden M, et al. An Outbreak of Human Coronavirus OC43 Infection and Serological Cross-reactivity with SARS Coronavirus. Can J Infect Dis Med Microbiol. Hindawi; 2006;17: 330–336. doi:10.1155/2006/152612

5. Hand J, Rose EB, Salinas A, Lu X, Sakthivel SK, Schneider E, et al. Severe Respiratory Illness Outbreak Associated with Human Coronavirus NL63 in a Long-Term Care Facility. Emerging Infectious Diseases. 2018;24: 1964–1966. doi:10.3201/eid2410.180862

6. Zeng Z-Q, Chen D-H, Tan W-P, Qiu S-Y, Xu D, Liang H-X, et al. Epidemiology and clinical characteristics of human coronaviruses OC43, 229E, NL63, and HKU1: a study of hospitalized children with acute respiratory tract infection in Guangzhou, China. Eur J Clin Microbiol Infect Dis. 2018;37: 363–369. doi:10.1007/s10096-017-3144-z

7. Killerby ME, Biggs HM, Haynes A, Dahl RM, Mustaquim D, Gerber SI, et al. Human coronavirus circulation in the United States 2014-2017. J Clin Virol. 2018;101: 52–56. doi:10.1016/j.jcv.2018.01.019

8. Pfefferle S, Oppong S, Drexler JF, Gloza-Rausch F, Ipsen A, Seebens A, et al. Distant relatives of severe acute respiratory syndrome coronavirus and close relatives of human coronavirus 229E in bats, Ghana. Emerging Infectious Diseases. 2009;15: 1377–1384. doi:10.3201/eid1509.090224

9. Huynh J, Li S, Yount B, Smith A, Sturges L, Olsen JC, et al. Evidence supporting a zoonotic origin of human coronavirus strain NL63. Journal of Virology. 5 ed. American Society for Microbiology Journals; 2012;86: 12816–12825. doi:10.1128/JVI.00906-12

10. Tao Y, Shi M, Chommanard C, Queen K, Zhang J, Markotter W, et al. Surveillance of Bat Coronaviruses in Kenya Identifies Relatives of Human Coronaviruses NL63 and 229E and Their Recombination History. Perlman S, editor. Journal of Virology. American Society for Microbiology Journals; 2017;91: 85. doi:10.1128/JVI.01953-16

11. Khalafalla AI, Lu X, Al-Mubarak AIA, Dalab AHS, Al-Busadah KAS, Erdman DD. MERS-CoV in Upper Respiratory Tract and Lungs of Dromedary Camels, Saudi Arabia, 2013–2014. Emerging Infectious Diseases. 2015;21: 1153–1158. doi:10.3201/eid2107.150070

12. Corman VM, Ithete NL, Richards LR, Schoeman MC, Preiser W, Drosten C, et al. Rooting the phylogenetic tree of middle East respiratory syndrome coronavirus by characterization of a conspecific virus from an African bat. Journal of Virology. American Society for Microbiology Journals; 2014;88: 11297–11303. doi:10.1128/JVI.01498-14

13. Corman VM, Eckerle I, Memish ZA, Liljander AM, Dijkman R, Jonsdottir H, et al. Link of a ubiquitous human coronavirus to dromedary camels. Proc Natl Acad Sci USA. National Academy of Sciences; 2016;113: 9864–9869. doi:10.1073/pnas.1604472113

14. Crossley BM, Mock RE, Callison SA, Hietala SK. Identification and characterization of a novel alpaca respiratory coronavirus most closely related to the human coronavirus 229E. Viruses. Multidisciplinary Digital Publishing Institute; 2012;4: 3689–3700. doi:10.3390/v4123689

15. Lau SKP, Woo PCY, Li KSM, Tsang AKL, Fan RYY, Luk HKH, et al. Discovery of a novel coronavirus, China Rattus coronavirus HKU24, from Norway rats supports the murine origin of Betacoronavirus 1 and has implications for the ancestor of Betacoronavirus lineage A. Sandri-Goldin RM, editor. Journal of Virology. American Society for Microbiology Journals; 2015;89: 3076–3092. doi:10.1128/JVI.02420-14

16. Wang W, Lin X-D, Guo W-P, Zhou R-H, Wang M-R, Wang C-Q, et al. Discovery, diversity and evolution of novel coronaviruses sampled from rodents in China. Virology. Academic Press; 2015;474: 19–27. doi:10.1016/j.virol.2014.10.017

17. Zhang J, Guy JS, Snijder EJ, Denniston DA, Timoney PJ, Balasuriya UBR. Genomic characterization of equine coronavirus. Virology. 2007;369: 92–104. doi:10.1016/j.virol.2007.06.035

18. Li K, Li H, Bi Z, Gu J, Gong W, Luo S, et al. Complete Genome Sequence of a Novel Swine Acute Diarrhea Syndrome Coronavirus, CH/FJWT/2018, Isolated in Fujian, China, in 2018. Matthijnssens J, editor. Microbiol Resour Announc. American Society for Microbiology; 2018;7: 466. doi:10.1128/MRA.01259-18

19. Zhou L, Li QN, Su JN, Chen GH, Wu ZX, Luo Y, et al. The re-emerging of SADS-CoV infection in pig herds in Southern China. Transbound Emerg Dis. John Wiley & Sons, Ltd; 2019;66: 2180–2183. doi:10.1111/tbed.13270

20. Zhou P, Fan H, Lan T, Yang X-L, Shi W-F, Zhang W, et al. Fatal swine acute diarrhoea syndrome caused by an HKU2-related coronavirus of bat origin. Nature. 1st ed. Nature Publishing Group UK; 2018;556: 255–258. doi:10.1038/s41586-018-0010-9

21. Yang Y-L, Qin P, Wang B, Liu Y, Xu G-H, Peng L, et al. Broad Cross-Species Infection of Cultured Cells by Bat HKU2-Related Swine Acute Diarrhea Syndrome Coronavirus and Identification of Its Replication in Murine Dendritic Cells In Vivo Highlight Its Potential for Diverse Interspecies Transmission. Gallagher T, editor. Journal of Virology. American Society for Microbiology Journals; 2019;93: 3134. doi:10.1128/JVI.01448-19

22. Edwards CE, Yount BL, Graham RL, Leist SR, Hou YJ, Dinnon KH, et al. Swine acute diarrhea syndrome coronavirus replication in primary human cells reveals potential susceptibility to infection. Proceedings of the National Academy of Sciences. National Academy of Sciences; 2020;4: 202001046. doi:10.1073/pnas.2001046117

23. Eckerle LD, Lu X, Sperry SM, Choi L, Denison MR. High fidelity of murine hepatitis virus replication is decreased in nsp14 exoribonuclease mutants. Journal of Virology. American Society for Microbiology Journals; 2007;81: 12135–12144. doi:10.1128/JVI.01296-07

24. Denison MR, Graham RL, Donaldson EF, Eckerle LD, Baric RS. Coronaviruses: an RNA proofreading machine regulates replication fidelity and diversity. RNA Biol. Taylor & Francis; 2011;8: 270–279. doi:10.4161/rna.8.2.15013

25. Smith EC, Denison MR. Implications of altered replication fidelity on the evolution and pathogenesis of coronaviruses. Curr Opin Virol. 2012;2: 519–524. doi:10.1016/j.coviro.2012.07.005

26. Graham RL, Baric RS. Recombination, Reservoirs, and the Modular Spike: Mechanisms of Coronavirus Cross-Species Transmission. Journal of Virology. 2010;84: 3134–3146.

27. Yang X-L, Hu B, Wang B, Wang M-N, Zhang Q, Zhang W, et al. Isolation and Characterization of a Novel Bat Coronavirus Closely Related to the Direct Progenitor of Severe Acute Respiratory Syndrome Coronavirus. Perlman S, editor. Journal of Virology. American Society for Microbiology; 2016;90: 3253–3256. doi:10.1128/JVI.02582-15

28. Hu B, Zeng L-P, Yang X-L, Ge X-Y, Zhang W, Li B, et al. Discovery of a rich gene pool of bat SARS-related coronaviruses provides new insights into the origin of SARS coronavirus. PLOS Pathogens. Public Library of Science; 2017;13: e1006698. doi:10.1371/journal.ppat.1006698

29. Kosakovsky Pond SL, Posada D, Gravenor MB, Woelk CH, Frost SDW. GARD: a genetic algorithm for recombination detection. Bioinformatics. 2006;22: 3096–3098.

30. Saberi A, Gulyaeva AA, Brubacher JL, Newmark PA, Gorbalenya AE. A planarian nidovirus expands the limits of RNA genome size. PLOS Pathogens. Public Library of Science; 2018;14: e1007314. doi:10.1371/journal.ppat.1007314

31. Debat HJ. Expanding the size limit of RNA viruses: Evidence of a novel divergent nidovirus in California sea hare, with a ~35.9 kb virus genome. 2018. doi:10.1101/307678

32. Fehr AR, Perlman S. Coronaviruses: an overview of their replication and pathogenesis. Methods Mol Biol. New York, NY: Springer New York; 2015;1282: 1–23. doi:10.1007/978-1-4939-2438-7_1

33. Coronaviridae Study Group of the International Committee on Taxonomy of Viruses. The species Severe acute respiratory syndrome-related coronavirus : classifying 2019-nCoV and naming it SARS-CoV-2. Nature Microbiology. Nature Publishing Group; 2020;5: 536–544. doi:10.1038/s41564-020-0695-z

34. Liu DX, Fung TS, Chong KK-L, Shukla A, Hilgenfeld R. Accessory proteins of SARS-CoV and other coronaviruses. Antiviral Research. 2014;109: 97–109. doi:10.1016/j.antiviral.2014.06.013

35. Di Tommaso P, Chatzou M, Floden EW, Barja PP, Palumbo E, Notredame C. Nextflow enables reproducible computational workflows. Nat Biotechnol. Nature Publishing Group; 2017;35: 316–319. doi:10.1038/nbt.3820

36. Katoh K, Misawa K, Kuma K-I, Miyata T. MAFFT: a novel method for rapid multiple sequence alignment based on fast Fourier transform. nar. Oxford University Press; 2002;30: 3059–3066.

37. Price MN, Dehal PS, Arkin AP. FastTree 2 – Approximately Maximum-Likelihood Trees for Large Alignments. Poon AFY, editor. PLoS ONE. Public Library of Science; 2010;5: e9490. doi:10.1371/journal.pone.0009490

38. Boni MF, Lemey P, Jiang X, Lam TT-Y, Perry BW, Castoe TA, et al. Evolutionary origins of the SARS-CoV-2 sarbecovirus lineage responsible for the COVID-19 pandemic. Nature Microbiology. Nature Publishing Group; 2020;382: 1–10. doi:10.1038/s41564-020-0771-4

39. Zhou H, Chen X, Hu T, Li J, Song H, Liu Y, et al. A Novel Bat Coronavirus Closely Related to SARS-CoV-2 Contains Natural Insertions at the S1/S2 Cleavage Site of the Spike Protein. Current Biology. Elsevier Ltd; 2020;: 1–12. doi:10.1016/j.cub.2020.05.023

40. Ge X-Y, Wang N, Zhang W, Hu B, Li B, Zhang Y-Z, et al. Coexistence of multiple coronaviruses in several bat colonies in an abandoned mineshaft. Virol Sin. Springer Singapore; 2016;31: 31–40. doi:10.1007/s12250-016-3713-9

41. Lam TT-Y, Jia N, Zhang Y-W, Shum MH-H, Jiang J-F, Zhu H-C, et al. Identifying SARS-CoV-2-related coronaviruses in Malayan pangolins. Nature. Nature Publishing Group; 2020;583: 282–285. doi:10.1038/s41586-020-2169-0

42. Xiao K, Zhai J, Feng Y, Zhou N, Zhang X, Zou J-J, et al. Isolation of SARS-CoV-2-related coronavirus from Malayan pangolins. Nature. Nature Publishing Group; 2020;583: 286–289. doi:10.1038/s41586-020-2313-x

43. Muth D, Corman VM, Roth H, Binger T, Dijkman R, Gottula LT, et al. Attenuation of replication by a 29 nucleotide deletion in SARS-coronavirus acquired during the early stages of human-to-human transmission. Sci Rep. Nature Publishing Group UK; 2018;8: 990–15177. doi:10.1038/s41598-018-33487-8

44. Su YCF, Anderson DE, Young BE, Linster M, Zhu F, Jayakumar J, et al. Discovery and Genomic Characterization of a 382-Nucleotide Deletion in ORF7b and ORF8 during the Early Evolution of SARS-CoV-2. Schultz-Cherry S, editor. mBio. 2020;11: e01610–20.

45. Pereira F. Evolutionary dynamics of the SARS-CoV-2 ORF8 accessory gene. Infect Genet Evol. Elsevier B.V; 2020;85: 104525–104525.

46. Young BE, Fong S-W, Chan Y-H, Mak T-M, Ang LW, Anderson DE, et al. Effects of a major deletion in the SARS-CoV-2 genome on the severity of infection and the inflammatory response: an observational cohort study. The Lancet. Elsevier; 2020;396: 603–611. doi:10.1016/S0140-6736(20)31757-8

47. Lau SKP, Feng Y, Chen H, Luk HKH, Yang W-H, Li KSM, et al. Severe Acute Respiratory Syndrome (SARS) Coronavirus ORF8 Protein Is Acquired from SARS-Related Coronavirus from Greater Horseshoe Bats through Recombination. Perlman S, editor. Journal of Virology. 2015;89: 10532–10547. doi:10.1128/JVI.01048-15

48. Vijgen L, Keyaerts E, Lemey P, Maes P, Van Reeth K, Nauwynck H, et al. Evolutionary history of the closely related group 2 coronaviruses: porcine hemagglutinating encephalomyelitis virus, bovine coronavirus, and human coronavirus OC43. Journal of Virology. 2006;80: 7270–7274. doi:10.1128/JVI.02675-05

49. Woo PCY, Lau SKP, Wernery U, Wong EYM, Tsang AKL, Johnson B, et al. Novel betacoronavirus in dromedaries of the Middle East, 2013. Emerging Infect Dis. 2014;20: 560–572. doi:10.3201/eid2004.131769

50. So RTY, Chu DKW, Miguel E, Perera RAPM, Oladipo JO, Fassi-Fihri O, et al. Diversity of Dromedary Camel Coronavirus HKU23 in African Camels Revealed Multiple Recombination Events among Closely Related Betacoronaviruses of the Subgenus Embecovirus. Pfeiffer JK, editor. Journal of Virology. American Society for Microbiology Journals; 2019;93: 255. doi:10.1128/JVI.01236-19

51. Lau SKP, Woo PCY, Yip CCY, Fan RYY, Huang Y, Wang M, et al. Isolation and characterization of a novel Betacoronavirus subgroup A coronavirus, rabbit coronavirus HKU14, from domestic rabbits. Journal of Virology. 6 ed. 2012;86: 5481–5496. doi:10.1128/JVI.06927-11

52. Nemoto M, Kanno T, Bannai H, Tsujimura K, Yamanaka T, Kokado H. Antibody response to equine coronavirus in horses inoculated with a bovine coronavirus vaccine. J Vet Med Sci. JAPANESE SOCIETY OF VETERINARY SCIENCE; 2017;79: 1889–1891. doi:10.1292/jvms.17-0414

53. Gong L, Li J, Zhou Q, Xu Z, Chen L, Zhang Y, et al. A New Bat-HKU2-like Coronavirus in Swine, China, 2017. Emerging Infectious Diseases. 2017;23: 1607–1609. doi:10.3201/eid2309.170915

54. Lau SKP, Woo PCY, Li KSM, Huang Y, Wang M, Lam CSF, et al. Complete genome sequence of bat coronavirus HKU2 from Chinese horseshoe bats revealed a much smaller spike gene with a different evolutionary lineage from the rest of the genome. Virology. Elsevier Inc; 2007;367: 428–439. doi:10.1016/j.virol.2007.06.009

55. Maganga GD, Pinto A, Mombo IM, Madjitobaye M, Mbeang Beyeme AM, Boundenga L, et al. Genetic diversity and ecology of coronaviruses hosted by cave-dwelling bats in Gabon. Sci Rep. Nature Publishing Group; 2020;10: 7314–13. doi:10.1038/s41598-020-64159-1

56. Vijgen L, Keyaerts E, Moës E, Thoelen I, Wollants E, Lemey P, et al. Complete genomic sequence of human coronavirus OC43: molecular clock analysis suggests a relatively recent zoonotic coronavirus transmission event. Journal of Virology. American Society for Microbiology Journals; 2005;79: 1595–1604. doi:10.1128/JVI.79.3.1595-1604.2005

57. Guo H, Hu B-J, Yang X-L, Zeng L-P, Li B, Ouyang S, et al. Evolutionary Arms Race between Virus and Host Drives Genetic Diversity in Bat Severe Acute Respiratory Syndrome-Related Coronavirus Spike Genes. Pfeiffer JK, editor. J Virol. 2020;94: e00902–20.

58. Li F. Structure, Function, and Evolution of Coronavirus Spike Proteins. Annu Rev Virol. Annual Reviews; 2016;3: 237–261. doi:10.1146/annurev-virology-110615-042301

59. Guindon S, Lethiec F, Duroux P, Gascuel O. PHYML Online—a web server for fast maximum likelihood-based phylogenetic inference. nar. 2005;33: W557–W559.

